# Killing by Type VI secretion drives clonal phase separation and the evolution of cooperation

**DOI:** 10.1101/063487

**Authors:** Luke McNally, Eryn Bernardy, Jacob Thomas, Arben Kalziqi, Jennifer Pentz, Sam Brown, Brian Hammer, Peter J. Yunker, William Ratcliff

**Affiliations:** Centre for Immunity, Infection and Evolution, School of Biological Sciences, University of Edinburgh, Edinburgh EH9 3FL, UK; Institute of Evolutionary Biology, School of Biological Sciences, University of Edinburgh, Edinburgh EH9 3FL, UK; School of Biology, Georgia Institute of Technology. Atlanta, USA. 30332.; School of Physics, Georgia Institute of Technology. Atlanta, GA, USA. 30332.

## Abstract

By nature of their small size, dense growth and frequent need for extracellular metabolism, microbes face persistent public goods dilemmas^1–5^. Spatial assortment can act as a general solution to social conflict by allowing extracellular goods to be utilized preferentially by productive genotypes^1,6,7^. Established mechanisms that generate microbial assortment depend on the availability of free space^8–14^; however, microbes often live in densely-packed environments, wherein these mechanisms are ineffective. Here, we describe a novel class of self-organized pattern formation that facilitates the development of spatial structure within densely-packed bacterial colonies. Contact-mediated killing through the Type VI secretion system (T6SS) drives high levels of assortment by precipitating phase separation, even in initially well-mixed populations that do not necessarily exhibit net growth. We examine these dynamics using three different classes of mathematical models and experiments with mutually antagonistic strains of *Vibrio cholerae* growing on solid media, and find that all appear to de-mix via the same ‘Model A’ universality class of order-disorder transition. We mathematically demonstrate that contact killing should favour the evolution of public goods cooperation, and empirically examine the relationship between T6SSs and potential cooperation through phylogenetic analysis. Across 26 genera of Proteobacteria and Bacteroidetes, the proportion of a strain’s genome that codes for potentially-exploitable secreted proteins increases significantly with both the number of Type 6 secretion systems and the number of T6SS effectors that it possesses. This work demonstrates how antagonistic traits—likely evolved for the purpose of killing competitors—can indirectly lead to the evolution of cooperation by driving genetic phase separation.

## Results and Discussion

Microbes are fundamentally social organisms. They often live in dense, surface-attached communities, and participate in a range of social behaviors mediated through the production and consumption of extracellular proteins and metabolites. Paradigmatic examples include the cooperative production of digestive enzymes^15^, metal chelators^16^, signaling molecules^15^, and the structural components of biofilms^17^. Many of these extracellular compounds are susceptible to social exploitation, in which non-producing ‘cheats’ gain an evolutionary advantage. If unchecked, this social exploitation can lead to the extinction of cooperative genotypes^18,19^. While it is widely recognized that the spatial segregation of cooperative microbes away from cheats can solve this cooperative dilemma^1,5,19^, relatively little work has investigated mechanisms that can generate genetic segregation within initially mixed communities.

Using a bacterial model system^20^ and mathematical modeling, we examine the biophysical basis of novel ecological structuring created by contact-mediated killing through the Type VI secretion system (T6SS). The T6SS is a potent mechanism of intermicrobial aggression, allowing bacteria to deliver lethal doses of effector proteins to adjacent competitors, while leaving clonemates with identical protective immunity proteins unscathed^21,22^.

**Figure 1.**
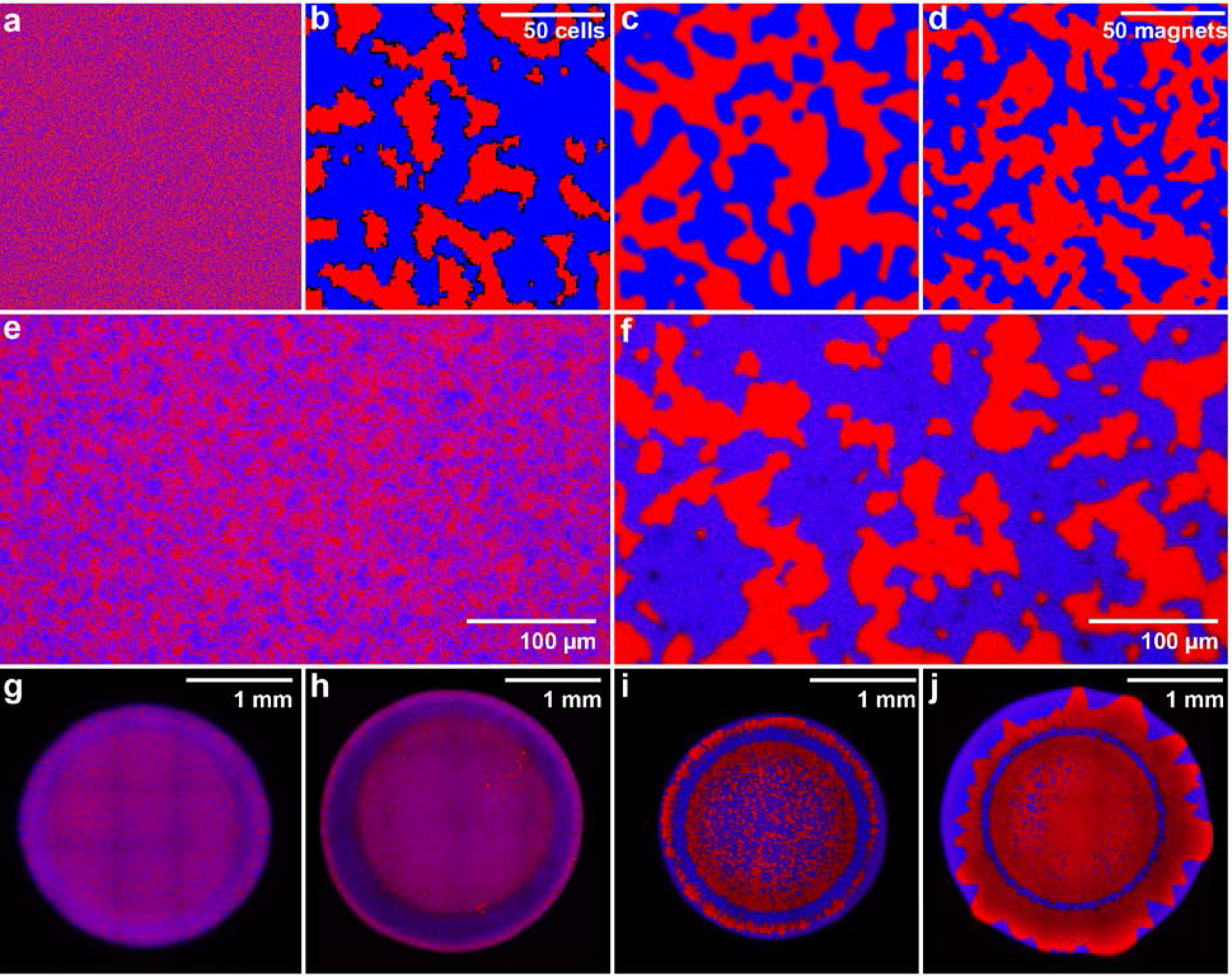
T6SS-mediated killing drives phase separation in dense bacterial populations. We modeled the dynamics of phase separation in fully-occupied, randomly seeded square lattices(a). Phase separation between red and blue bacteria capable of mutual killing occurred in an individual based model (b), in a partial differential equation model (c) and in an Ising spin model (d). No phase separation occurred between red (C6706) and blue (692-79)T6SS^-^ mutants of *Vibrio cholerae* (*ΔvasK;* e), in contrast to T6SS^+^ strains (f). We varied the efficacy ofT6SS while still allowing for growth by culturing *V. cholerae* at a range of temperatures-17°C (h), 25°C (i) and 30°C (j). T6SS^-^ controls cultured at 25°C did not phase separate (g).

Our system illustrates the profound effect of T6SS-mediated killing on emergent spatial patterning of a surface attached population. Mathematical modeling suggests that an initiallywell-mixed population of mutual killers should rapidly undergo phase separation due to ‘selfish herd’ dynamics^23^, as the cells within genetically-uniform groups no longer risk T6SS-mediated death. Indeed, we observe rapid phase separation in three distinct classes of models, all starting with a randomly seeded population on a 2D lattice (Fig. 1a): an Individual Based Model (IBM; Fig. 1b; Supplementary Movie 1), a Partial Differential Equation model (PDE; Fig. 1c; Supplementary Movie 2), and the classic Ising spin model^24^ (Fig. 1d; Supplementary Movie 3). Similarly, initially-well mixed populations of two *Vibrio cholerae* strains (C6706 and 692-79; Table S1) capable of mutual T6SS-mediated killing (Fig. S1) underwent phase separation (Fig. 1f, i & j). Non-killing controls (*ΔvasK, i.e.*, T6SS^-^; Fig. S1) and T6SS^+^ mutual killers cultured at low temperatures which impede T6SS activity^25^ remained well-mixed (Fig. 1e, g & h).

**Figure 2.**
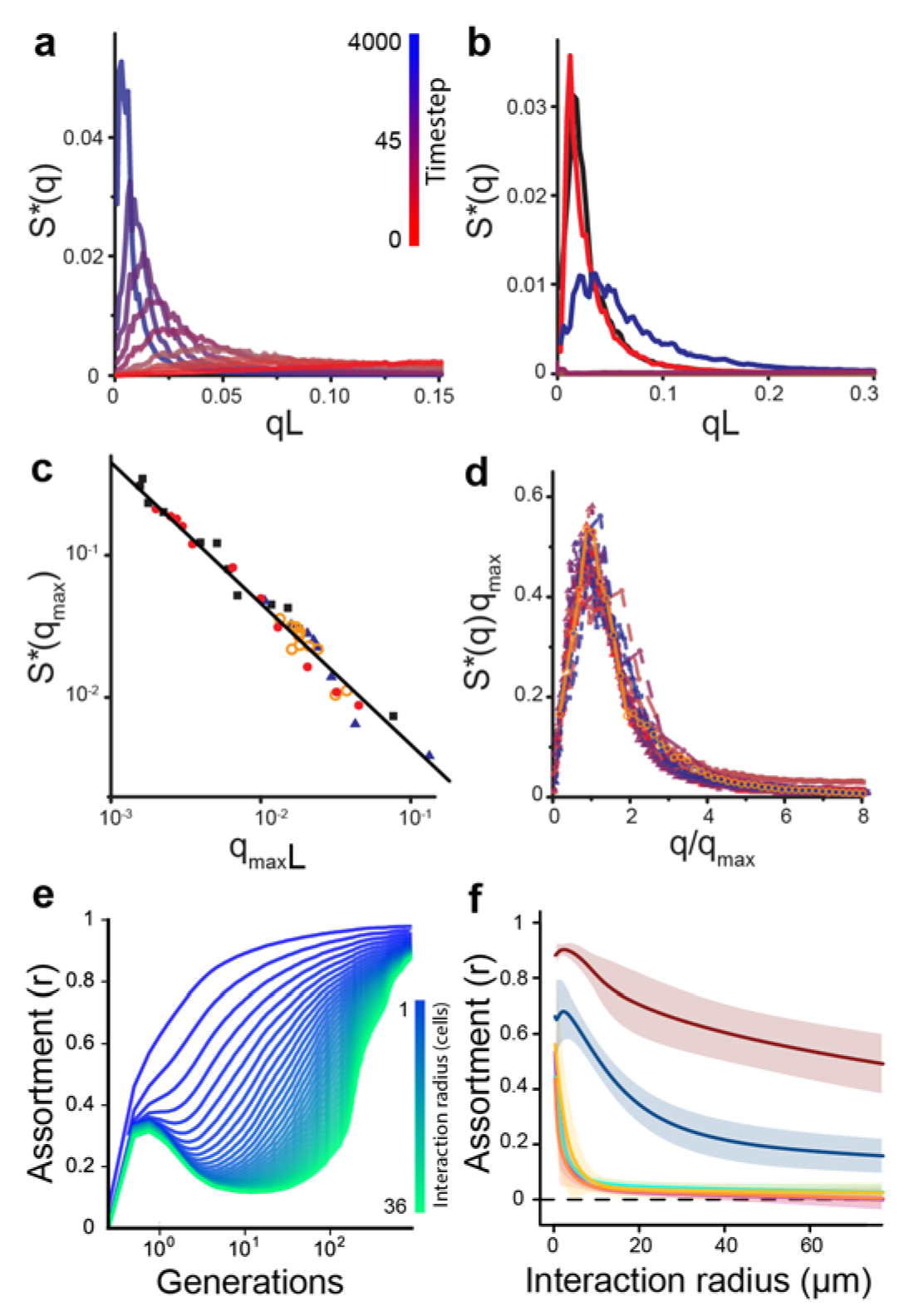
Structural analysis of models and experiments. The normalized static structure factor *S*(q)*, plotted vs. wavenumber *q* multiplied by cell size *L* for the Individual Based Model (IBM; a) and for experiments (b). In the latter, the red and black lines depict two separate fields of view of *V*. *cholera*e strains C6706 and 692-79, started at an initial ratio of 1:6, while blue indicates a 1:8 inoculation ratio. The brown line depicts T6SS^-^ mutants, and purple indicates mutual killers grown at 17°C for 24 h (all others grown at 25°C). Mutual killing drives phase separation, increasing the peak in *S*(q)* at smaller values of *q_max_*. The relationship between *S\**(*q_max_*) and *q_max_* is summarized in (c) with 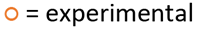 data (25°C and a 1:6 inoculation ratio, as in panel b), 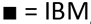, 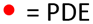 model (d=0.01), and 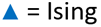 model (T=1); all three models and the experiments follow a universal 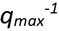 trend. *S*(q)* curves collapse when *S*(q)q_max_* is plotted versus *q*/*q_max_* (d), indicating that all models and experiments are undergoing the same coarsening process. Colour denotes model timestep, as in (a), while symbols indicate type of model or experiment, as in (c). We also examine the creation of spatial structure by calculating a biological metric, assortment (r), through time in the IBM (e) and after 24 h in experiments (f). Mutual killers were grown at 30°C (red), 25°C (blue) and 17°C (green). Defective killers were grown at 30°C (purple), 25°C(teal) and 17°C (orange). Plotted is the mean assortment of ≥ 3 replicate populations ± 95% confidence intervals.

To determine whether our models and experiments undergo the same type of order-disorder transition, we quantitatively examined the dynamics of phase separation in each. We first computed the Fourier-transformed structure factor, *S(q)*; to facilitate comparisons between models and experiments, we calculate 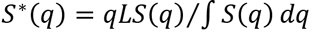 (Fig. 2a & b), where *L* is the size of a unit cell. The peak in *S*(q)* identifies the most common characteristic length scale (the inverse of q) of clonal groups, and the height of the peak is related to how often it occurs in the lattice (*i.e.* the strength of patterning at that length scale). At early timesteps (Fig. 2a), or for non-killing controls (Fig. 2b), *S*(q)* is relatively flat, as expected for a well-mixed population lacking a characteristic length scale. T6SS-mediated killing causes a peak to appear in *S*(q)*, which grows in height and moves to smaller values of *q* (longer length scales) as the population grows increasingly structured. This progression of *S*(q)* is a hallmark of phase separation^26^. The location of the first peak in *S*(q)*, denoted *q_max_*, scales as 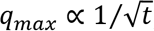, while the height of the peak scales^27^ as 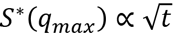.It is ambiguous how to relate simulation time to experimental time; instead, we plot *S*(q_max_)* versus *q_max_*. Remarkably, all models (IBM, PDE and Ising) and experiments fall on the same line 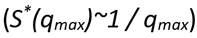 (Fig. 2c), a relationship consistent with the “Model A” order-disorder phase separation process^28^ described by the Allen-Cahn equation 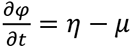, which relates the change in local concentration, *φ*, over time to diffusion and the chemical potential, *μ* and stochastic fluctuations (Supplementary Equations 2)^29^. Cellular mobility has a surprising effect on phase separation: rather than impeding phase separation, it accelerates it by enhancing killing at the borders of clonal patches(Supplementary Equations 1, Supplementary Figure 3 and Supplementary Movie 4).

To provide biological context for this process of phase separation, we calculated clonal assortment (*r*), for the IBM (Fig. 2e) and the *Vibrio* experiments (Fig. 2f). T6SS-mediated killing resulted in the creation of highly structured populations with high assortment over long length scales (Fig. 2e&f), which can protect diffusible public goods from consumption by competing strains^30,31^. To explore the effect of T6SS-mediated killing on the evolutionary stability of public goods cooperation, we introduced a diffusible cooperative good into our PDE model. We considered two competing strains: a cooperator that secretes an exoproduct into its environment at an individual cost, and a non-producing cheat that, all else equal, grows faster than the cooperator as it does not pay the cost of production. In this model, cellular growth rates for both strains depend on the local concentration of the diffusible exoproduct. We find that T6SS-mediated killing protects cooperation in two different ways. In a non-spatial (*i.e.*, constantly mixed) environment, T6SS-mediated killing can allow cooperators to resist invasion by rare cheats owing to the cooperators’ numerical dominance in antagonistic interactions (Figure 3c).However, in a spatially-defined environment, phase separation driven by T6SS-mediated killing physically separates producers from cheats, expanding the conditions favouring cooperation and allowing them to invade a population of cheats from rarity (Figure 3d&e and Supplementary Movie 5; see proof in Supplementary Equations 3).

**Figure 3.**
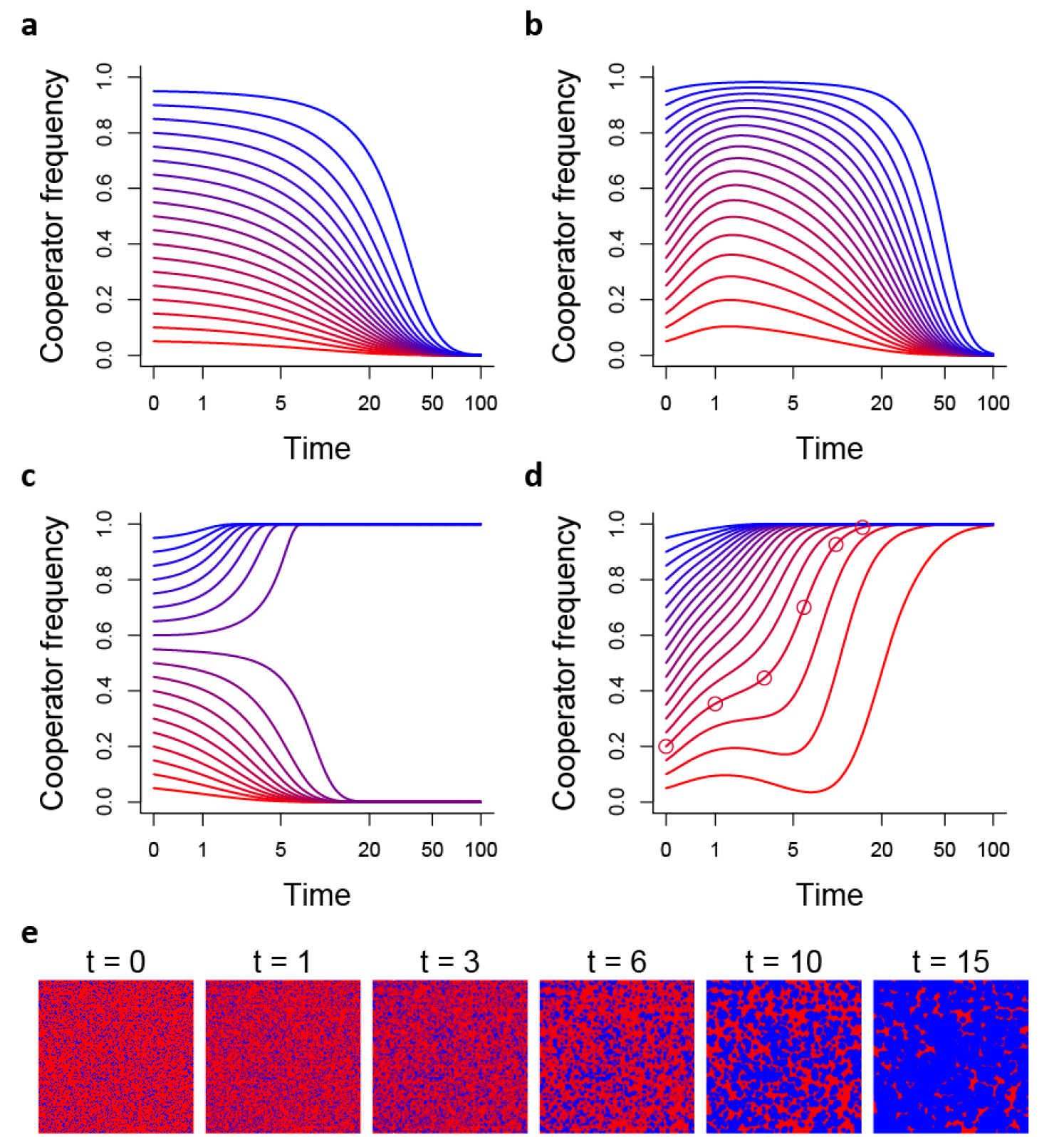
Phase separation favours the evolution of cooperation. The dynamics of competition between cooperators and cheats are shown through time for different starting frequencies. In the absence of T6SS-mediated killing, cooperation is not favoured in either a well-mixed environment (a) or a spatially-defined environment (b). In a non-spatial environment with killing via T6SS, cooperators can be protected from cheats when common owing to their advantage in antagonistic interactions, but cannot invade from rarity (c). In contrast, the high assortment created by phase separation allows cooperators to invade from rarity and spread to fixation (d). The spatial organization of cooperators (blue) and cheats (red) during competition is shown in (e). Panels correspond to the time-points marked by circles in (d).

Our models and experiments demonstrate that T6SS-mediated killing can rapidly structure surface-attached bacterial populations, generating favourable conditions for the evolution ofpublic-goods cooperation^1,7,10,32^. Does T6SS-mediated killing have a similar effect in the real world, where ephemeral resources, physical disturbance, and intense competition may impede phase separation? We approach this question phylogenetically, examining the relationship between the proportion of each genome coding for potentially-exploitable extracellular products and its T6SS complexity, with the rationale that a greater number of T6SSs and effector proteins should generally allow for the exclusion of a greater diversity of competitors. We constructed a Bayesian phylogenetic mixed model of T6SS-containing Proteobacteria and Bacteroidetes (Fig. 4a) using 439 genomes from 26 genera. Secretome size is positively correlated with both the number of T6SSs (Fig. 4b, d, Table S2) and T6SS effector proteins (Fig. 4c, e, Table S2) present. These results are also robust in univariate analyses (Tables S3 and S4) and to the inclusion of genome size as a predictor (Table S5). As our analyses include many closely related strains (*e.g.*, many *Helicobacter pylori*, Fig. 4a), most (91%) of the variance in secretome size is explained by the phylogenetic relationships among strains. Nonetheless, the number of T6 secretion systems and T6SS effectors are important predictors of secretome size, explaining 8% of the total, and 90% of the non-phylogenetic variance in secretome size. While, as with any phylogenetic analysis, alternative hypotheses cannot be ruled out entirely, these results strongly suggest that T6SS-mediated killing creates conditions that favour exoproduct evolution across a broad diversity of bacterial taxa.

Phase separation is well-known to drive pattern formation in biology^14^, but has mainly been investigated using either Turing activator-inhibitor feedbacks^33,34^, or positive density-dependent movement, described by the Cahn-Hilliard equation^14,35–37^. In this paper we describe a third general mechanism of self-organized pattern formation: targeted killing of non-kin competitors. This drives a ‘Model A’ phase separation; the kinetics of this coarsening process—described by the Allen-Cahn equation—only depend on a few cellular details. While we explore this process in bacteria, it is probably more general, applying to other organisms that kill adjacent non-kin (*e.g.*, allelopathy in plants^38^ and animals^39^).

**Figure 4.**
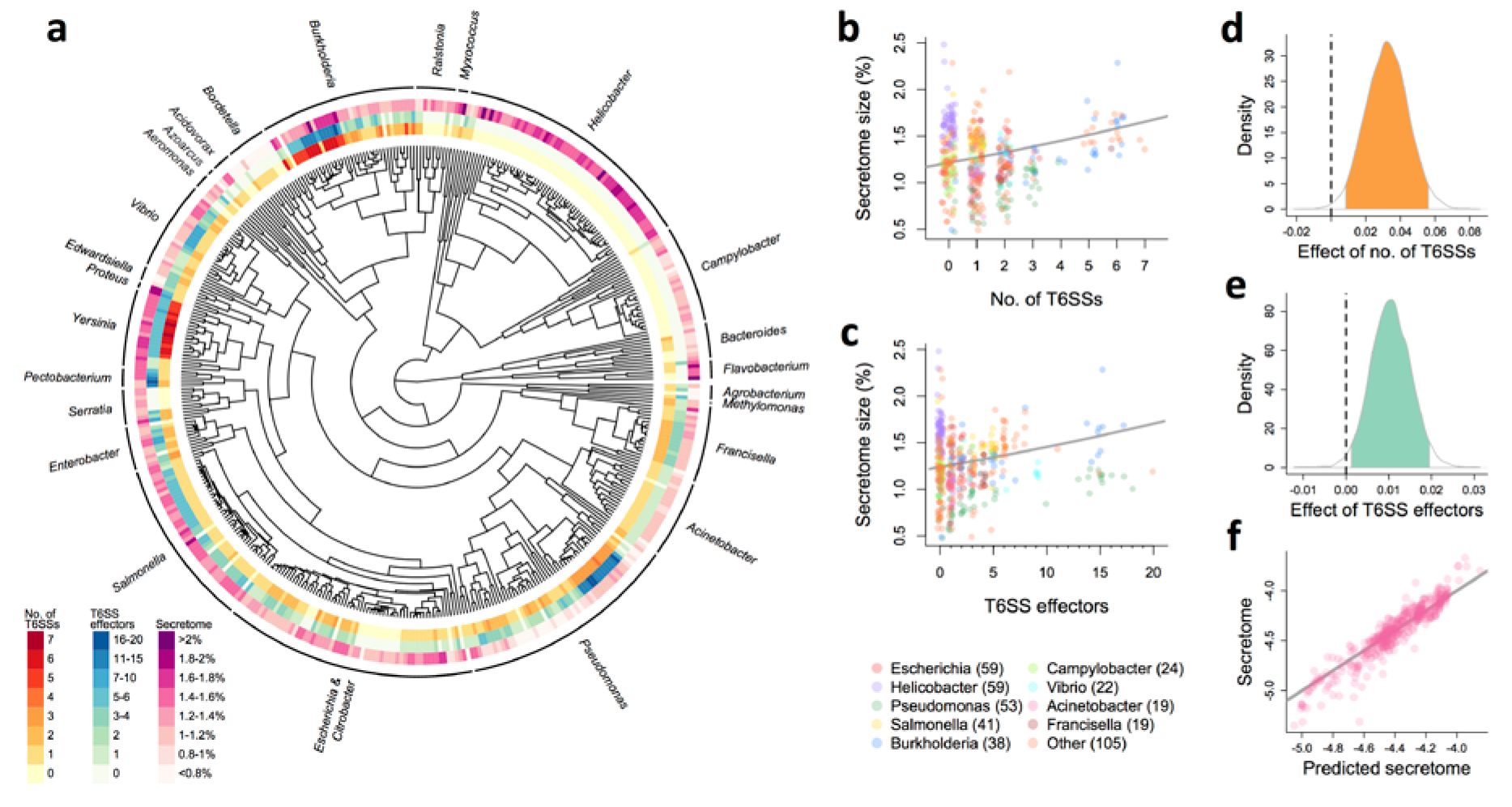
T6SS is associated with investment in other extracellular metabolites across Proteobacteria and Bacteroidetes. The phylogenetic distribution of T6SS, T6SS effectors, and secretome size across the Proteobacteria and Bacteroidetes (a). Secretome size of a strain (expressed as a percentage of genome size) increases with both its number of T6SSs (b) and T6SS effectors (c). Lines are the fits of univariate BPMMs (Tables S3 and S4). Posterior distributions of the effects of the numbers of T6SS (d) and T6SS effectors (e) on secretome size from the multivariate BPMM (Table S2). 95% credible intervals of the estimates are shaded. Plot of observed against predicted secretome size from the multivariate BPMM (f), including effects of the number of T6SS, number of T6SS effectors and phylogeny. The line represents a 1:1 mapping.

In recent years, there has been a growing appreciation that many microbial behaviors requiring extracellular metabolism are susceptible to social exploitation. Here we show how simple cell-cell aggression can, as a consequence, create a structured population favourable to cooperation. Because T6SSs are common (found in ∼25% of gram negative bacteria^21^), and microbes often live in dense communities, phase-separation driven by contact-mediated killing may play a fundamental role in defining the genetic composition and ecosystem-level functionality of microbial communities globally.

## Acknowledgements

This work was supported by NSF grant #DEB-1456652 to W.C.R. and NASA Exobiology grant #NNX15AR33G to W.C.R.; by the Gordon and Betty Moore Foundation grant #4308.07 to B.K.H., NSF grant #MCB-1149925 to B.K.H. and Wellcome Trust grant #WT095831 to S.B. L.M. was supported by HFSP grant #RGP0011/2014 to S.B. We would like to thank David Yanni, Jonathan Michel, Shane Jacobeen, and Wouter Ellenbroek for helpful comments.

### Author contributions

All authors actively participated in the planning of this research. E.B., J.T., and B.H. constructed the *Vibrio* mutants and performed all bacterial experiments. E.B., J.T., P.J.Y. and A.K. performed and analyzed the microscopy. L.M. and P.J.Y. wrote and analyzed the PDE model. L.M. performed the phylogenetic analysis. W.C.R. wrote the individual based model and script to calculate assortment, P.J.Y. and A.K. wrote the Ising spin model and script to calculate S(q). W.C.R., L.M., E.B. and P.J.Y. wrote the first draft of the paper, all authors contributed to revision.

**Methods.** The full methods section is attached as a supplemental document.

The **Supplementary Information** includes four figures, five supplementary movies, five tables, three mathematical appendices, a dataset, and a phylogeny.

